# Implicit food odour priming effects on reactivity and inhibitory control towards foods

**DOI:** 10.1101/2020.01.27.920967

**Authors:** Marine Mas, Marie-Claude Brindisi, Claire Chabanet, Stéphanie Chambaron

## Abstract

The food environment can interact with cognitive processing and influence eating behaviour. Our objective was to characterize the impact of implicit olfactory priming on inhibitory control towards food, in groups with different weight status. Ninety-two adults completed a modified Affective Shifting Task: they had to detect target stimuli and ignore distractor stimuli while being primed with non-attentively perceived odours. We measured reactivity and inhibitory control towards food pictures. Priming effects were observed on reactivity: participants with overweight and obesity were slower when primed with pear and pound cake odour respectively. Common inhibitory control patterns toward foods were observed between groups. We suggest that non-attentively perceived food cues influence bottom-up processing by activating distinguished mental representations according to weight status. Also, our data show that cognitive load influences inhibitory control toward foods. Those results contribute to understanding how the environment can influence eating behaviour in individuals with obesity.

## Introduction

Studies have shown that individuals with obesity tend to have poorer inhibition capacities when it comes to food [1,2]. In our food-abundant environment, this tendency inevitably leads to overeating, i.e. eating more than one’s physiological needs. This type of impaired inhibition can naturally lead to weight gain and even to obesity.

The combination of excess calorie intake and a lack of caloric expenditure results in weight excess, overweight, and often obesity. This phenomenon is related to our environment: for most people in modern-day society, food is abundant and easily accessible. Moreover, daily exercise is now a choice rather than an obligation. Scientists have therefore introduced the idea of the “obesogenic” environment, inferring that the influence of the environment is a key feature of the current obesity epidemic. According to Swinburn et al., “the physiology of energy balance is proximally determined by behaviours and distally by environments” [3]. However, it is still difficult to explain how, why, and under which conditions the obesogenic environment can influence food choices on an information-processing level. Indeed, obesity has a multifactorial aetiology, and researchers have highlighted genetic, metabolic, social, psychological, cognitive, and environmental factors that contribute to the maintenance and development of obesity [3–6].

Independently of their surroundings, people are, by nature, attracted to food [7]. Indeed, foraging for nutritious food is one of the key roles of the brain functions as food is essential for survival [8]. The brain preferentially directs its limited resources toward energy-dense food stimuli [8,9]. Food also induces reward in the form of pleasure in the dopaminergic pathways of the brain, which is similar to the cognitive processing of addictive substance cues [10,11]. Those two aspects provide a solid base to establish that food stimuli are salient in the environment [12]: they are more prone to visually attract attention, and consequently, undergo quickly cognitive processing, which affects decision-making. [13]. In an obesogenic environment, biased decision-making in favour of high energy-dense food choices inevitably leads to weight gain.

To study the modulation of behaviour by food cues, several studies focused on using priming [14,15]. Priming is how incidental stimuli (stimuli that are perceived without the individual’s awareness or appreciation of their influence) are shown to influence higher-order cognitive and behavioural outcomes [16]. Incidental stimuli that alter human food behaviour can be visual (advertisements, [15]), auditive [17], but also olfactive. Indeed, olfaction is strongly tied to food intake as food odours generally signal food availability [18]. Several studies have shown that food odour priming might modify several aspects of eating behaviour, such as attitudes to foods [19], food choices [20], food intake [21,22], and bottom-up cognitive processing of food stimuli [23].

In a previous study, we highlighted the differing influence of incidental olfactory food cues on the stimulus-driven^1^ cognitive processing of food pictures in individuals with different weight statuses [23]. Indeed, when primed with non-attentively perceived odours signalling high energy-dense (HED) foods, participants with obesity tended to show greater orienting attentional biases (*i.e. the individual tendency to automatically orient one’s attention toward specific stimuli*) toward food pictures than when primed with non-attentively perceived odours signalling low energy-dense (LED) foods. This tendency was reversed for individuals with normal weight status, and different from the pattern of attentional orienting toward foods in individuals with overweight. In sum, implicit olfactory priming with food odours can either increase or decrease the perceptual salience^2^ of foods in different ways according to weight status by influencing the bottom-up processing of such stimuli. We consequently wondered whether olfactory priming with food cues could also have differentiated effects on goal-directed or “top-down” processes such as inhibitory control. This contribution would help us to clarify the links between the processing of food cues and food-related decision-making.

Inhibitory control is part of the executive functions, which are cognitive functions responsible for transmission between endogenous (mood, thoughts, sensations) and exogenous (environmental) events. Executive functions are involved in problem-solving and decision-making, which are necessary for the execution of goal-directed actions [24–26]. Inhibitory control is a remarkable executive function that makes it possible for us to stay consistent with our behavioural intentions on attentional, cognitive and behavioural levels. Many researchers have conceptualized several theoretical models of its structure [27,28]. According to Friedman and Miyake (2004), there are three defined components of inhibitory control: (a) attentional control, allowing us to focus our attention on stimuli of interest and to avoid wasting mental resources on non-pertinent stimuli, (b) cognitive inhibition, namely the ability to resist proactive interference from prepotent stimuli in information processing, and (c) self-control, the ability to control one’s behaviour instead of acting impulsively [27]. Each of these three components is involved in a specific type of stimulus processing, which helps individuals to adapt to changing situations by enabling voluntary behaviours and inhibiting possible perturbations.

The hypothesis of a deficit in inhibitory control among individuals with obesity has been widely explored by researchers in an effort to explain why weight loss remains difficult, and to find innovative opportunities to reduce obesity [10]. Such a deficit could lead to a decrease in the ability to pursue goal-directed behaviour, such as maintaining a healthy lifestyle. In this line of study, some authors showed that individuals with obesity have lower inhibitory control, [2,25,29,30] while other studies found no differences related to weight status [31,32]. No consensus has been found so far, potentially due to the diversity of methodologies [33]. Additionally, other variables (such as frequent comorbidities in obesity, or specific eating styles) are susceptible to modulate inhibitory control capacities beyond weight status [32,34–36]. Applied to food-choice behaviour, low inhibitory control is related to excessive consumption of HED foods, especially in contexts of consumption facilitation [37,38]. Moreover, in an obesogenic context where there is an overload of information, few cognitive resources remain available to inhibit one’s attention, thoughts and behaviours. This may guide individuals toward default choices, namely palatable but unhealthy foods [7].

Some sensory cues create a context of facilitation by guiding the individual toward consumption [39] while offering opportunities to succumb to the temptation of palatable foods. Among these cues, food odours have a strong influence; they signal the availability of foods without necessarily raising awareness [40,41]. To our knowledge, our study is the first to explore the relationship between a context of facilitation and inhibitory control toward foods (high and low energy-dense foods vs. neutral non-food stimuli) in male and female adults of various weight statuses (normal-weight, overweight, obese) and with no eating disorder. New data on how food stimuli modulate cognitive processing might help to understand how individuals are influenced by our obesogenic environment. Moreover, the lack of inhibitory control toward foods is one problematic aspect of eating behaviour. Disentangling its mechanisms could explain some health-deleterious food choices in obesity.

The first aim of this study was to characterize inhibitory control toward food pictures in individuals with normal-weight, overweight and obesity. Our second aim was to study how olfactory priming affected top-down processes in individuals with various weight statuses, by measuring their inhibitory control capacities when non-attentively exposed to olfactory food cues compared to non-exposed. Our main hypothesis was that, compared with neutral stimuli (objects), individuals facing food stimuli would have decreased inhibitory control, especially when the food stimuli were HED. We expected that this deficit would be increased in individuals with higher weight status, especially when non-attentively primed with olfactory food cues.

## Material and methods

### Participants

One hundred and twenty-four adults aged from 20 to 60 years old were recruited and grouped according to their body mass index (BMI, kg/m^2^, [42,43]; 38 individuals with obesity (OB), 45 individuals with overweight (OW), and 41 individuals with normal weight (NW). Participants were recruited from the population registered in the Chemosens Platform’s PanelSens database. This database complies with national data protection rules and has been vetted by the appropriate authorities (Commission Nationale Informatique et Libertés - CNIL). Participants were contacted by an e-mail from the platform which invited them to respond to a questionnaire investigating inclusion and exclusion criteria mentioned below. The study was conducted in accordance with the Declaration of Helsinki and was approved by the Comité d’Evaluation Ethique de l’Inserm (CEEI, File number IRB 0000388817-417). This research study adhered to all applicable institutional and governmental regulations concerning the ethical use of human volunteers.

Exclusion criteria were: age under 18 or over 60 years old, diagnosis of a chronic disease (such as type 2 diabetes, cardiovascular disease, or hypertension), regular medical treatment causing cognitive impairment (antipsychotic, anxiolytic, or antidepressant), olfactory impairment (anosmia, hyposmia, chronic sinusitis) and a history of bariatric surgery. Additionally, participants who were sick (cold or flu symptoms) at the time of the experiment were asked to postpone their appointment with the laboratory in order to ensure that they did not have an impaired sense of smell during the session.

Written informed consent was obtained from participants before their participation, though they came to the session under a false pretence (i.e., to participate to a computerized experiment on picture categorization). At the end of the experiment, participants were entirely debriefed and told the real purpose of the study. In return for their participation, the participants received a €10 voucher at the end of the session

### Measurements

#### An adaptation of the Affective Shifting Task

In order to measure inhibition toward foods, we adapted the affective shifting task [44,45] modified by Mobbs, Iglesias, Golay, & Van der Linden, 2011. This task is based on the Go/No-go paradigm (for a review, see Gomez, Ratcliff, & Perea, 2007). In this task, participants must both (a) detect target stimuli (go trials) by pressing the spacebar on a computer keyboard and (b) withhold their response to distracter stimuli (no-go trials). Participants were instructed to respond as fast and as accurately as they could. During the task, two instruction types alternated: target stimuli were either food stimuli (“food set”, HED or LED food pictures) or objects (“object set”, tools or household objects). Stimuli were selected from FoodPics [48] and rigorously paired in terms of perceptual and consumer properties according to the procedure used in [23].

The task comprised 3 blocks of 112 trials each. Each block comprised 4 sets (order: food-object-food-object) of 28 trials each (28% HED trials, 28% LED trials and 44% objects trials, in a pseudo-random order without three pictures of the same type appearing consecutively). See fig 1. for details. Each set began with oral instructions about the target stimuli (food or object) given through a headset, then a fixation cross appeared for 500ms at the centre of a black screen. Subsequently, pictures appeared one by one for 500ms, with an inter-stimuli-interval of 900ms consisting of a white fixation cross on a black screen that participants were instructed to fixate. Commission and omission errors were signalled to the participant by a short sound conveyed by the headset. Blocks were separated by 1-minute pauses during which experimenters took the headsets off participants and invited them to relax. Prior to measurements, participants completed a brief training session comprising 4 sets of 10 trials in order to familiarize them with the task. They were asked to rate their hunger level on a 10-point Likert scale before and after the modified Affective Shifting Task.

**Fig 1.**
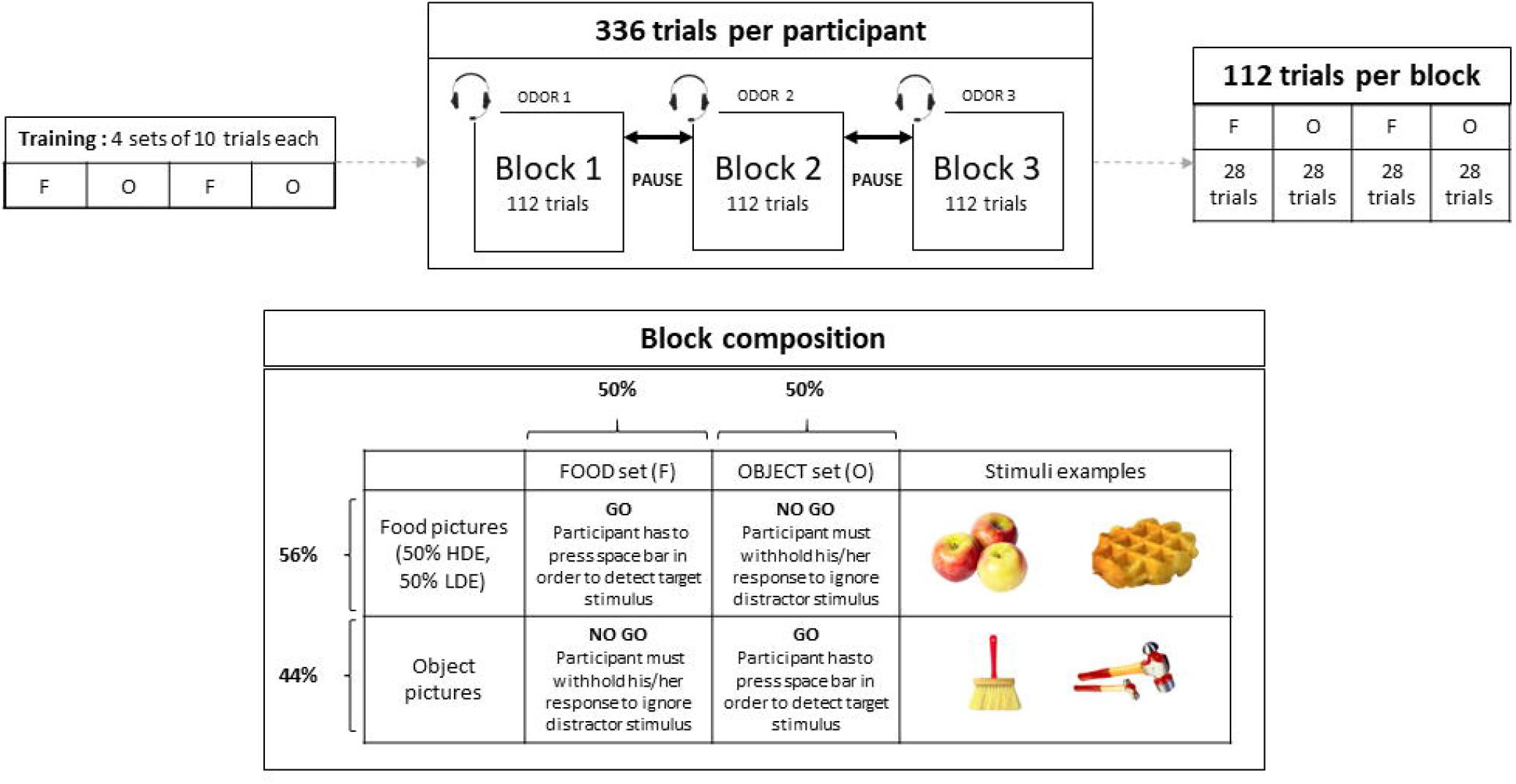
Composition of blocks, sets and trials of the modified Affective Shifting Task. F = food, O = object.

For each subject and for each experimental trial, we collected the reaction times (RT), the presence of a commission error (detecting a distractor stimulus) and the presence of an omission error (not detecting a target stimulus). Reaction times corresponded to the time between the appearance of the stimulus on screen and the moment the participant pressed the space bar to detect it (0 to 500ms). Commission errors corresponded to situations in the no-go trials in which the participant pressed the space bar, indicating a lack of response inhibition to distractor stimuli. Omission errors corresponded to go-trials for which the participant did not press the space bar to detect the target stimulus, indicating a lack of attention to the given stimulus [45,49].

#### Priming

In order to non-attentively expose participants to olfactory food cues, we used the olfactory priming paradigm developed by Marty & al. in 2017 [19,23]. In this paradigm, participants perform three identical blocks of a computerized task (here, the modified Affective Shifting Task) while wearing a headset with a microphone. The headsets are used to provide instructions to participants, and, unbeknownst to participants, the microphones are used as brackets for odorized microphone foams. Task blocks are separated by short pauses during which experimenters discreetly switch the headsets in order to non-attentively expose participants to different olfactory food cues through the odorized foams of the headset’s microphone. Our study had three different olfactory priming conditions: odour signalling HED foods (fatty sweet pound cake odour), odour signalling LED foods (fruity pear odour) and control condition in which the foam was not odorized.

Participants come to the laboratory under a false pretence (here, taking part in a study on picture categorization) so they do not guess the presence of olfactory cues during the session. At the end of the three blocks of the task, participants complete an investigation questionnaire in which they have to guess the aim of the experiment and indicate whether they noticed anything particular during the task that could have influenced their performance. Participants mentioning odours or headsets in this questionnaire are excluded from the study. This step ensures that no odour or headset change was perceived, which allows the implicit quality of the priming [23].

#### Global Cognitive Capacities

To ensure that differences in cognitive performance during the modified Affective Shifting Task were not due to general cognitive deficits, participants performed standardized tests, namely the Go/No-go and flexibility subtests of the computerized Test of Attentional Performance (TAP) neuropsychological test battery [50].

The Go/no-Go subtest explores response inhibition through a simple task in which the participant must detect target stimuli “X” and withhold a response when presented with distractor stimuli “+”. The flexibility subtest assesses shifting abilities in mental flexibility. In this subtest, two stimuli appear, one on the left and one on the right side of the screen. One of the stimuli is round while the other is an angular shape. The participant must detect whether the round shape is on the left or on the right side of the screen by pressing the corresponding key with the dominant hand through several trials. Participants were given a brief training before each subtest. The assessment began systematically with the Go/No-go subtest.

### Session

Participants came to the laboratory at 12 p.m. They were instructed to refrain from eating, drinking anything except water, wearing scented cosmetics, smoking or chewing gum for 3 hours prior to the session. They began the session with the three blocks of modified Affective Shifting Task, followed by the investigation questionnaire and a hunger rating on a 10-point Likert scale. Then, they were administered the two subtests of the TAP [50], namely Go/No-go and Flexibility, in order to check their global cognitive performance. Afterwards, participants filled a computerized version of the Questionnaire for Eating Disorder Diagnosis – Q-EDD [51,52] in order to identify and exclude participants with potential eating disorders. Finally, participants passed the European Test for Olfactory Capacities – ETOC [53] in order to ensure that they could correctly detect and identify odours. At the end of the session, the weight and height of each participant were measured, individually, in a separate room by the experimenter.

### Data preparation

Since instruction shifts modulate task difficulty [54], so we created a two-level covariate to account for the cognitive load generated by the change of instructions between tasks (food-object-food-object). The two levels were “CL+” for the first 14 trials of each set and “CL-” for the second 14 trials of each set (total of 28 trials). The CL+ condition refers to a situation in which the individual becomes familiar with new instructions (detecting foods in food sets and detecting objects in object sets) and the implementation of the instructions is automatized during the set. In the CL-condition, the individual is already familiar with the instructions, implicating a lower cognitive load. This two-level covariate was integrated in further linear mixed models that are described below.

During data preparation, reaction times (RTs) inferior to 150ms were excluded from analysis because they reflect stimulus anticipation [46]. In order to analyse global reaction speed, we summarized, for each participant, RTs for which the spacebar was pressed (go trials without omissions and no-go trials with errors) by using the median per condition (olfactory prime type x stimulus type x cognitive load). For errors, we calculated the proportion of errors on no-go trials for each participant in each condition (olfactory prime type x stimulus type x cognitive load). For omission errors, the proportion of omissions among the go trials per condition was calculated for each participant.

For each dependent variable (RTs, proportion of commission errors, proportion of omission errors), we estimated a linear mixed model. The model initially involved four fixed factors (weight status group x stimulus type x olfactory prime type x cognitive load), all interactions, and the individual as a random factor. We also added covariates such as age, global cognitive performance (flexibility and Go/no-Go) and sex. We then simplified the model by removing non-significant terms except if they were involved in a significant higher-order term. Contrasts were used to interpret significant main effects and interactions.

Statistical analysis was performed with R.3.4.3 software [55] using linear mixed models (nlme package v. 3.1-131 [56]) to explain reactivity to stimuli expressed in median RTs, inhibitory control deficit expressed in proportion of errors, and inattention expressed in proportion of omissions. Specific contrasts were subsequently tested using the contrast package [57,58]. The significance threshold was set at 0.05. Full data are available in the Supporting Information Files, see S1 Dataset.

## Results

### Sample characteristics

At the end of the tests, 33 participants were excluded from the sample (see details in Fig 2). Indeed, 25 declared that they had smelled an odour during the session, meaning that the priming was not implicit for those participants. Five participants were screened as disordered eaters according to the Q-EDD, and two more participants were excluded because their answers to the ETOC indicated that they had low olfactory capacities (hyposmia or anosmia). One participant was excluded from analysis because data from the flexibility subtest were missing.

**Fig 2.**
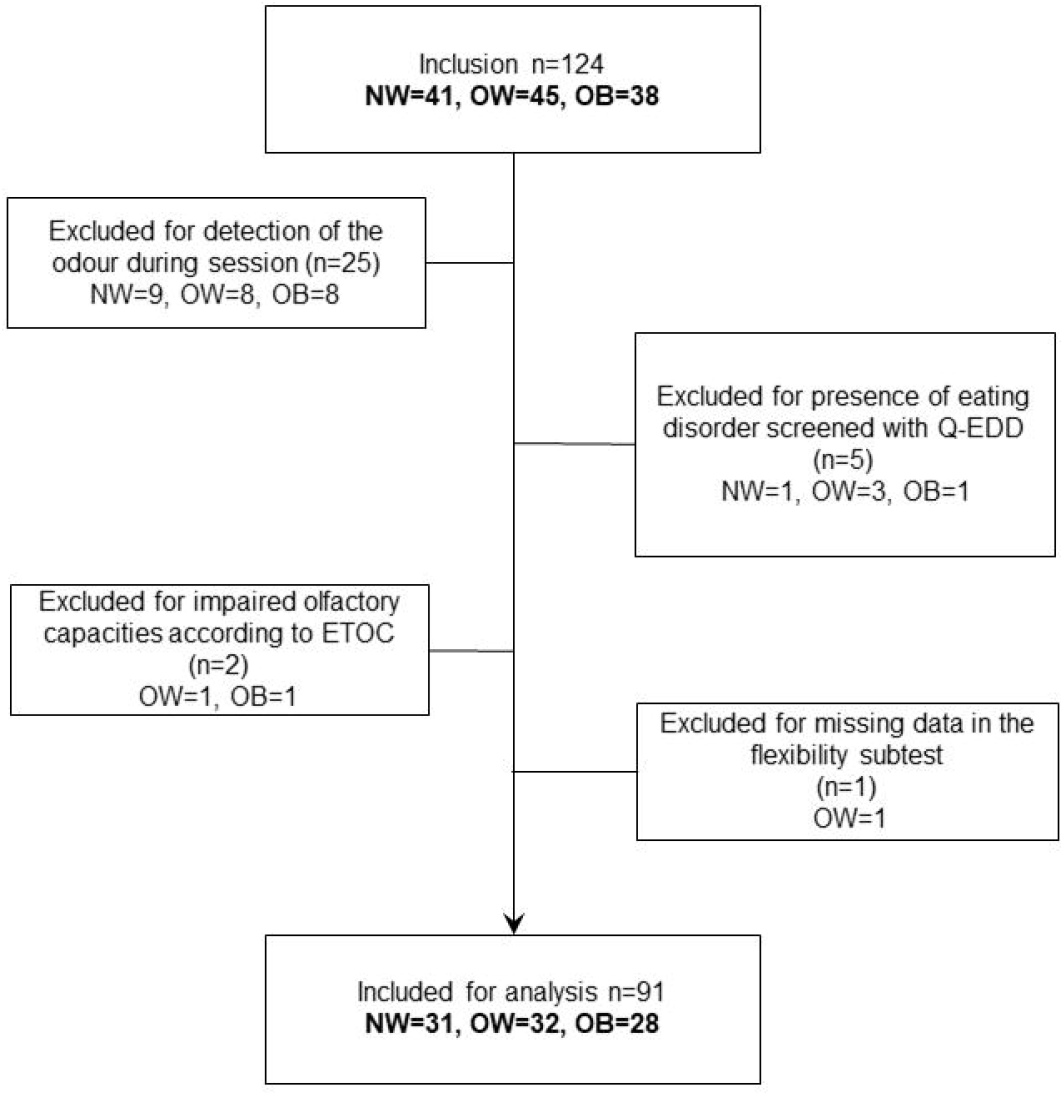
Flowchart of exclusions. NW = participants with normal weight, OW = participants with overweight, OB = participants with obesity.

Finally, 91 participants remained eligible for analysis: 31 participants with normal-weight, 32 participants with overweight and 28 participants with obesity (according to their BMI measurements).

When comparing the sociodemographic data of the 3 BMI groups, ANOVA test were used for quantitative variables and Chi2 tests were used for categorical variables (sex ratio, educational level). No significant differences were observed in age, sex ratio, educational level, hunger level before the session or variations in hunger during the session. To measure the change in hunger, the hunger level before the session was subtracted from hunger level after session (both had been rated on a 10-point Likert scale before and after the modified Affective Shifting Task).

For the scores on the TAP sub-tests, performances are indicated in T-scores for the number of errors (reflective of inhibitory control capacities) in the Go/No-go subtest. For the flexibility subtest, a global performance index (GPI), [50] was calculated for each participant, based on the T-scores for reaction times and the T-scores concerning the number of errors for each participant (0.707 * (T_Median RT_ + T_Number of errors_ – 100)). If the GPI is positive (>0), individual performance is interpreted as being above the mean performance of the reference sample (normative data), while if it is negative (< 0), it is interpreted as being lower than the average performance of the reference sample (normative data). T-scores are normalized scores based on the percentile of scores in a reference population (mean=50, SD=10, [50]). Average performance is comprised between 43 and 57 (corresponding to the 25 and 75 percentiles, respectively) and T-scores are adjusted on sex, gender and educational level. No significant difference in global inhibition (Go/No-go) and flexibility were found between weight status groups. Details of sociodemographic characteristics are displayed in Table 1.

**Table 1:** Participants characteristics.

|  | Weight status |  |  |  |  |  |
| --- | --- | --- | --- | --- | --- | --- |
|  | Normal-weight (NW) |  | Overweight |  | Obesity |  |
|  | n=31 (34%) |  | (OW) |  | (O) |  |
|  |  |  | n=32 (35%) |  | n=28 (31%) |  |
|  | Mean | (SD) | Mean | (SD) | Mean | (SD) |
| **Age (y): p=0.70** | 43.41 | (11.07) | 44.15 | (8.76) | 41.89 | (11.30) |
| **BMI (kg/m<sup>2</sup>): p<0.001** | 21.95<sup>a</sup> | (1.77) | 27.28<sup>b</sup> | (1.37) | 36.43<sup>c</sup> | (5.75) |
| **Hunger level before session (1-10): p=0.19** | 6.33 | (2.14) | 5.62 | (2.75) | 5.07 | (2.97) |
| **Variation in hunger: p=0.60** | 0.45 | (0.75) | 0.75 | (1.42) | 0.44 | (1.82) |
| **TAP Go/No-go subtest – (T-score): p=0.15** | 48.45 | (6.65) | 46.71 | (6.03) | 44.67 | (9.22) |
| **TAP Flexibility subtest – (GPI): p=0.92** | 1.41 | (6.09) | 1.96 | (8.60) | 1.26 | (7.45) |
|  | n | % | n | % | n | % |
| **Sex: p=0.69** |  |  |  |  |  |  |
| **Women** | 19 | (61%) | 16 | (50%) | 17 | (61%) |
| **Men** | 12 | (39%) | 16 | (40%) | 11 | (39%) |
| **Level of education: p=0.85** |  |  |  |  |  |  |
| **< 14 years** | 16 | (52%) | 16 | (50%) | 16 | (57%) |
| **> 14 years** | 15 | (48%) | 16 | (50%) | 12 | (43%) |

Quantitative variables expressed as mean (SD)
<sup>a, b, c</sup> Superscript letters are associated with values (means or numbers), same letters indicating that the difference
between values is not significant. P values indicate the significance of the weight status effect. GPI = Global
Performance Index.

### Reaction times

The main effect of the type of stimulus [F_(2, 1536)_=46.94, p<0.001)], the interaction between weight status and olfactory prime type [F_(4, 1536)_= 3.21, p=0.012] and the interaction between weight status and cognitive load [F_(2,1536)_]=5.47, p=0.004] reached significance in the RT linear mixed model. Age and global cognitive performance (Flexibility and Go/no-Go performances) were also kept in the model as they were significant as covariates. Main results are shown in Fig 3.

**Fig 3.**
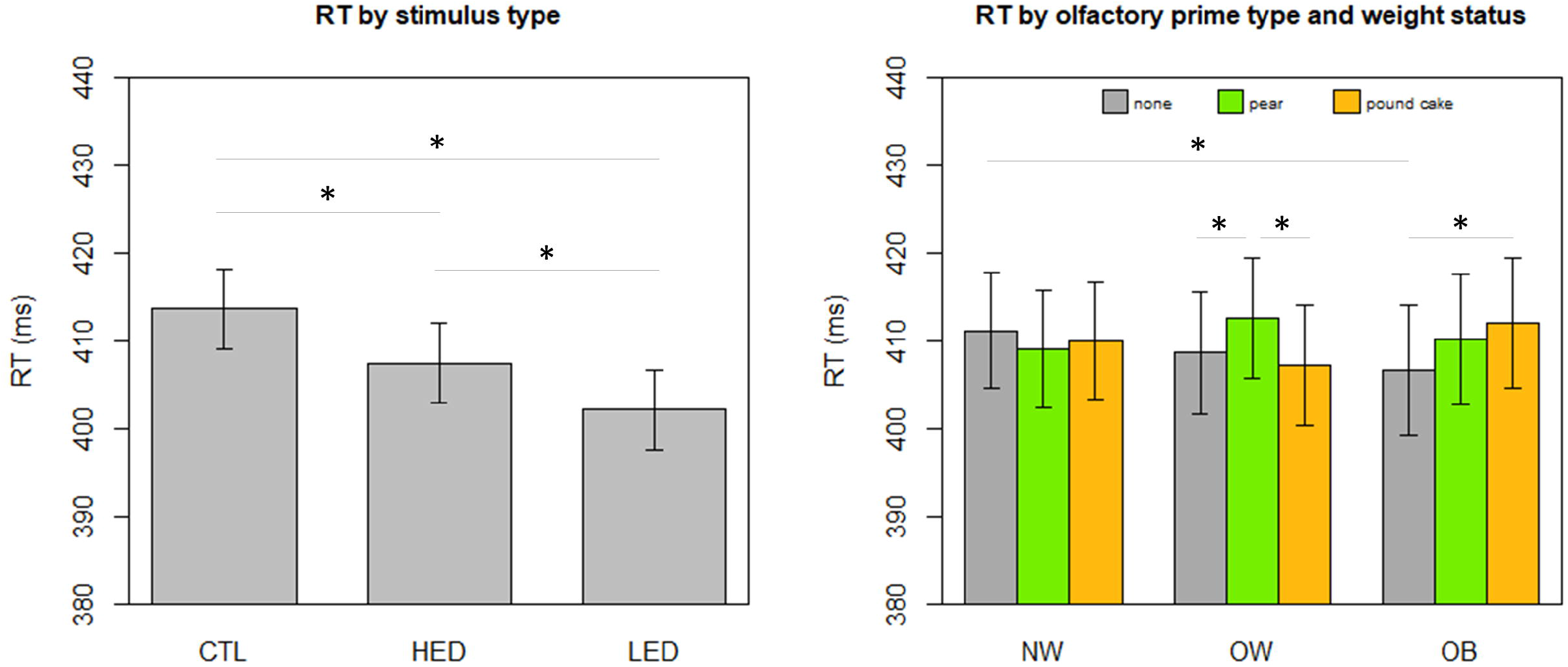
(left) RT by stimulus type, CTL=objects (control) pictures, HED=high energy-dense foods pictures, LED=low energy-dense foods pictures, averaged on olfactory prime type, cognitive load condition and weight status. (right) RT by olfactory prime type and weight status (NW=normal-weight, OW=overweight, OB=obesity), averaged on stimulus type and cognitive load condition. Predicted values and 95% confidence intervals. * p < 0.05.

Regarding the main effect of stimulus type, individuals detected food pictures faster than object pictures [HED vs objects = −6.20ms (p<0.001), LED vs objects = −11.53ms (p<0.001)], and responded quicker to LED food pictures than HED food pictures [LED vs HED = −5.33ms, (p<0.001)].

Regarding the interaction between weight status and olfactory prime type, participants with obesity were slower to detect stimuli of all types when primed with a pound cake odour [OB, pound cake odour vs none=+5.30ms, (p=0.01) and, non-significantly, when primed with a pear odour [OB, pear vs. none=+3.54ms, (p=0.09)]. Participants with overweight were slower to detect stimuli when primed with a pear odour [OW, pear vs pound cake=+5.33ms (p=0.01)] and non significantly, when they were primed with a pear odour vs. no odour [OW, pear vs none=+3.95ms, (p=0.049)]. On the contrary, participants with normal weight showed no significant difference between RT when primed with a pound cake odour (p=0.58) or with a pear odour (p=0.30).

When we looked at the interaction between weight status and cognitive load, only normal-weight individuals had different reaction times depending on cognitive load conditions. More specifically, they were slower when the cognitive load was higher [NW, CL+ vs CL-=+5.22ms, (p=0.002)]. In addition, in the higher cognitive load conditions, normal-weight participants tended to be slower than participants with overweight [CL+, NW vs. OW=+8.34ms, (p=0.07)]. However, these results only approached significance.

### Proportion of commission errors

Three terms of the commission errors linear mixed model reached significance: the main effect of stimulus type [F_(2, 1542)_=51.36, p<0.001], the main effect of cognitive load condition [F_(1,1542)_=24.29, p<0.001] and the interaction between cognitive load and stimulus type [F_(2, 1542)_= 5.29, p=0.005]. Sex and global cognitive performance on the Go/no-Go subtest were also kept in the model as they were significant as covariates. Results are shown in Fig 4.

**Fig 4.**
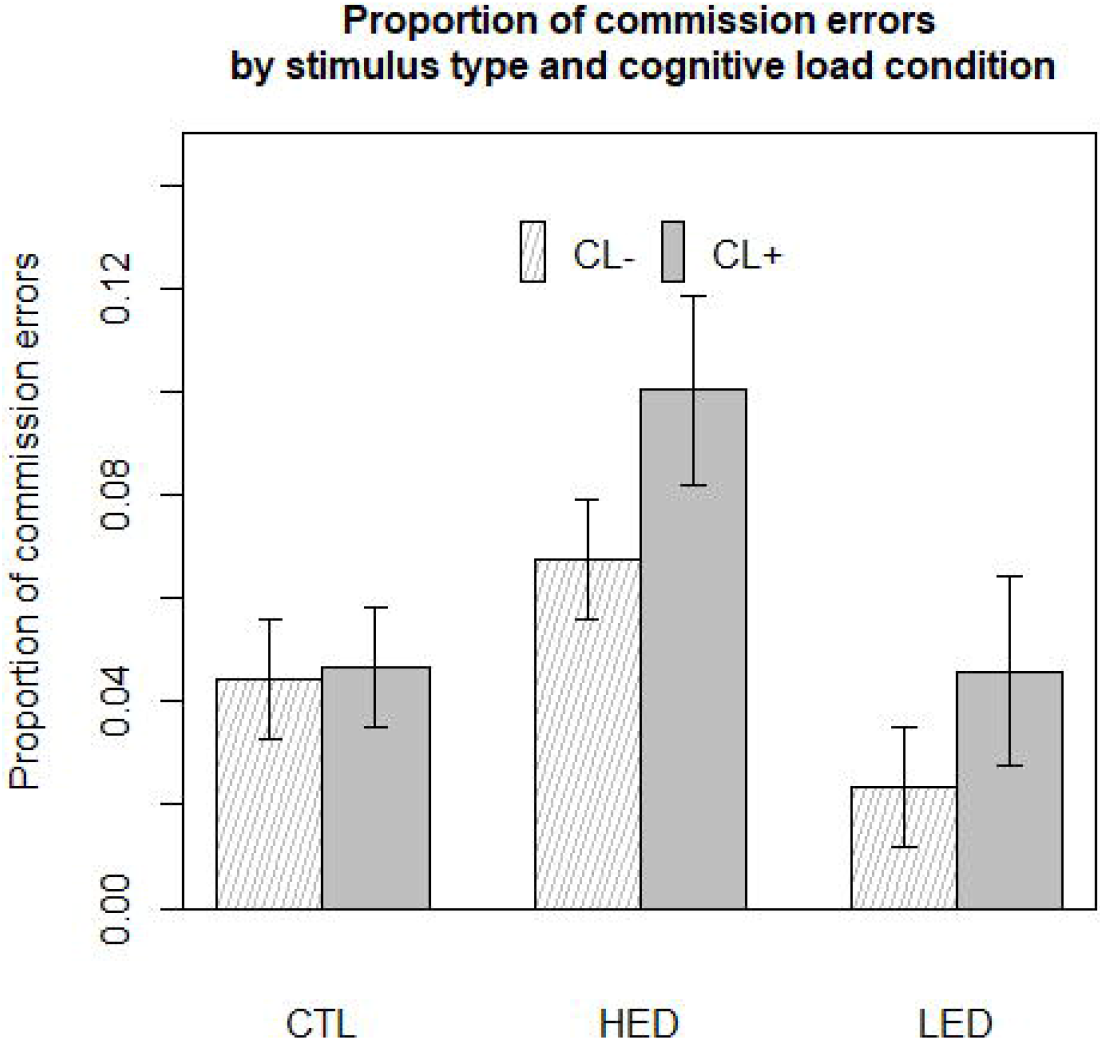
Proportion of commission errors by stimulus type and cognitive load condition averaged on olfactory prime type and weight status. CL+=high cognitive load condition, CL-=low cognitive load condition, CTL=objects (control) pictures, HED=high energy-dense food pictures, LED=low energy-dense food pictures. Predicted values and 95% confidence intervals.

Concerning the effect of cognitive load, participants made 43% more commission errors in the CL+ condition than in the CL-condition. [CL+ vs. CL- = +2.07 errors, p<0.001].

Stimulus type effect was dependent on cognitive load condition. In both the high and low cognitive conditions, participants made on average 84% more commission errors when facing HED food stimuli than when facing objects [HED vs objects=+3.95 errors, p<0.001]. Participants also made 142% more commission errors when facing HED food stimuli than when facing LED food stimuli [HED vs. LED=+4.92 errors, p<0.0001]. A slight difference in the amount of commission errors made was observed between LED food stimuli and objects, but it did not reach significance in the CL+ condition [CL+, LED vs. objects= −1 error, (NS, p=0.059)]. Nevertheless, in the CL-condition, participants made 90% more commission errors for objects than for LED food stimuli [CL-objects vs LED=+2.09 errors, (p=0.004)].

Participants made more commission errors in CL+ conditions than in CL-conditions for food stimuli: 48% and 96% more commission errors were made in the CL+ condition for HED and LED food stimuli, respectively [HED, CL+ vs. CL-= +3.50 errors, p <0.001; LED, CL+ vs. CL-= +2.48 errors, p<0.001)]. Participants did not make a significantly different proportion of commission errors between high and low cognitive load conditions when facing object stimuli [objects, CL+ vs. CL-=+0.22 errors, p=0.75].

In sum, HED food pictures induced more disinhibition than LED food and object pictures. The cognitive load modulated this disinhibition for food stimuli but not for neutral stimuli.

### Proportion of omission errors

Only two terms of the linear mixed model reached significance for the proportion of omission errors: main effect of type of stimulus [F_(2,1541)_=90.45, p<0.001] and interaction of weight status group and type of stimulus [F_(4,1541)_=2.67, p=0.03]. Age and global cognitive performance on the Go/no-Go subtest were also kept in the model as they were significant as covariates.

Concerning the main effects of stimulus type, participants made 117.2% more omission errors when facing HED food stimuli than facing LED food stimuli [HED vs. LED=+6.35 omissions errors, p<0.001]. They also made significantly fewer omission errors for food stimuli than for objects: 15.6% and 61.2% less omission errors were made for HED and LED food stimuli, respectively, in comparison with object stimuli [HED vs. objects=-3.49 omission errors, p<0.0001, LED vs. objects= −9.85 omission errors, p<0.001].

When we focused on the interaction between weight status group and stimulus type, we found that NW participants made more omissions than OW participants when facing HED food stimuli [HED, NW vs. OW=+4.32 omission errors, (p=0.044)]. No other effects approached significance. In sum, food pictures, especially HED foods, elicited more omission errors than neutral pictures in all participants.

## Discussion

Our objective was to characterize deficits in inhibitory control toward foods in different weight status groups (NW, OW, OB), and to assess the impact of implicit olfactory priming (pound cake, pear, control) on such processes.

### Global performance

Global performance for inhibitory control was similar for all groups in our sample, as measured by the Go/no-Go subtest from the TAP, and for flexibility as measured with the flexibility subtest from the same battery. Contrary to previous findings [29,30,59–61], inhibitory control and mental flexibility capacities were similar regardless of weight status. In addition, the number of commission errors, omission errors and reaction times in the modified Affective Shifting Task revealed no significant differences according to weight status when participants were not primed with a non-attentively perceived food cue. This suggests that common processes in the detection of stimuli and inhibition capacities are not dependent on weight status.

### Reactivity to foods

In our experiment, all participants reacted more quickly to food pictures than to neutral pictures. This highlights that food stimuli undergo faster processing, which is in line with previous literature [23,62–66]. Indeed, food is essential for survival (i.e. a primary motivated goal of the individual) and has a rewarding quality, which are characteristics of a salient stimulus [12]. So food stimuli appear to be processed more quickly, which explains the increased reactivity to foods in all individuals.

The present study distinguished the approach bias for low energy-dense (LED) foods and for high energy-dense (HED) foods. While comparing RTs for high-calorie and low-calorie foods, Meule et al. suggested that longer RTs for HED foods indicated increased attention toward them. This relates to the fact that HED foods capture attention more forcefully than LED foods in the early stages of cognitive processing, which is consistent with our previous work on orienting attentional biases [23]. Moreover, it seems that HED food stimuli tend to capture attentional focus for longer periods of time than LED food stimuli. This might be behaviourally reflected in reaction times, as highlighted by neuroimaging studies showing discriminative patterns of activity in the brain for high and low-calorie food stimuli [67,68]. In our experiment, individuals were faster to detect LED food stimuli than HED food stimuli. This finding may relate to the attentional dimension of inhibitory control [27], which could be impaired by the perception of HED food pictures.

HED food stimuli processing might initially be facilitated by the high perceptual salience of high-calorie foods. We suggest that over time, the detection of HED food stimuli is impaired by their capacity to attract the focus of attention (slowed disengagement, [69,70]), which slows behavioural responses. On the contrary, LED food stimuli processing might be facilitated by the earlier identification of fruit stimuli in our experiment. As food stimuli, LED stimuli are also salient. However, their processing is not impaired by the attentional approach bias elicited by the higher appetitive quality of HED food stimuli. This effect results in a decrease in reaction times for LED foods compared with HED foods, partly explaining why participants had shorter RT and fewer omission errors for LED food stimuli than for HED food stimuli in our experiment.

### Differences in vulnerability to food cues in individuals with higher weight status

Concerning global reaction times, we found some priming effects for individuals with overweight and obesity. More specifically, individuals with obesity and with overweight were slower to detect all kinds of stimuli when primed with a pound cake odour and a pear odour, respectively, regardless of the go/no-go instructions. In our study, the odour signalling HED or LED foods could have slowed the bottom-up processing of foods by adding another element to take into account in the detection of stimuli. This indicates that olfactory food cues were implicated in the detection process by slowing RT in individuals of higher weight status. We consequently hypothesize that implicit priming effects only influence the bottom-up processing of food cues.

The result of the priming effect seen here is congruent with the results of previous studies [19,23]. In an earlier study, we found that implicit priming of olfactory food cues had differentiated effects: individuals with obesity were more vulnerable to a non-attentively perceived pound cake odour in their bottom-up processing of food cues [23]. For individuals with overweight in the present study, the effect of the pear odour is consistent with a study by Marty & al [19] in which olfactory pear and pound cake primes had differentiated effects when they were non-attentively perceived by children with overweight. Indeed, these children were more prone to choose fruit in a forced-choice task when they were non-attentively primed with a pear odour. The authors explained this result by hypothesizing that individuals with overweight might be more confronted to the idea of “dieting” in their daily lives, and so this concept might be more easily activated by a non-attentively perceived odour signalling a LED food. Future research could focus on understanding why odours signalling LED foods seem to affect individuals with overweight while odours signalling HED foods affect individuals with obesity. These food types may differentially activate certain concepts and mental representations in individuals according to weight status.

### Inhibitory control toward foods

Though we hypothesized that individuals with higher weight status would show less inhibitory control toward foods than lean individuals, it was not the case in our experiment. In fact, we found common patterns of inhibitory control toward food stimuli in individuals across the weight status spectrum.

In our experiment, participants made more commission errors when they were facing HED food stimuli. No difference was found in regard to weight status, which is congruent with part of the literature [71,72]. This observation strongly suggests that the lack of inhibition toward foods is a common process for all individuals and it is also consistent with the idea that the rewarding quality of HED foods makes them more appealing [73–75], leading to an increased approach bias. The salience of HED foods combined with the associated approach bias makes the detection of HED food stimuli a prepotent response for the individual. A prepotent response is cognitively more difficult to inhibit than other response options, which need to be inhibited in order to exhibit goal-congruent behaviour. This effect appears to be even stronger when cognitive load is high because individuals make significantly more commission errors toward HED food stimuli in this condition.

We found different patterns of inhibitory control toward HED and LED foods, indicating that the top-down processing of those stimuli is differentiated. In lower cognitive load conditions, individuals made fewer commission errors when facing LED food stimuli than when facing HED food or object stimuli. We can thus presume that fruits (LED foods) are processed faster than other stimuli. This assumption is supported by the work of Leleu et al., 2016 [76], who showed that fruit pictures elicited earlier event-related responses in the brain than other food types (vegetables, HED foods) during a food discrimination task.

### Priming effects: why does implicit priming only impact bottom-up processes?

In our study, we tested whether implicit priming with olfactory food cues would impact inhibitory control, a decision-driven, or “top-down” process measured by the proportion of commission errors made by participants in each olfactory condition. Unexpectedly, no priming effect was observed for commission errors, contrary to the effects observed with the exact same olfactory priming paradigm used in a Visual Probe Task to measure orienting attentional biases (a stimulus-driven, bottom-up process) [23]. Because orienting attentional biases are data-driven processes, sensory inputs are important determiners of behavioural response in such tasks [77]. Moreover, the Visual Probe Task needed less top-down cognitive effort than the modified Affective Shifting Task. Hoffman-Hensel & al, 2017, who observed that cognitive effort altered the neural processing of food odours, found that involvement in multiple tasks decreased participants’ perception of odour intensity [78]. We can suppose that implicit olfactory cues effects may not have been strong enough to act as a facilitation context impairing the inhibitory control.

According to the I-RISA model of addictive behaviour, rewarding cues have an increased salience for individuals, which is related to lower inhibitory control in cognitive tasks and associated with specific activations in the brain [11,79,80]. In our study, participants showed similar patterns of inhibitory control toward foods, but the implicit priming of olfactory food cues (supposed to act as a facilitation context and increase the salience of visual food stimuli) did not seem to modulate inhibitory control performance. Some works from the field of addictions and neuroscience suggest that implicit cues (such as masked cues or subliminal pictures) might activate different areas of the brain (limbic system) than explicit rewarding cues (prefrontal cortex) in addicted individuals [79,81,82]. This assumption supports the fact that implicit and explicit priming might differentially modulate bottom-up and top-down processing of stimuli. An alternative explanation for the absence of priming effect on inhibitory control in our study might be that implicit olfactory stimuli might only target the bottom-up processing of food pictures.

### Modulation of inhibitory control capacities toward food by cognitive load

Finally, the high cognitive load condition induced slower reaction times and more commission errors for all participants facing all types of stimuli in each olfactory condition. This reflects the worse performance and higher mental effort required to complete the task [83] and confirms that the first half of each set was more difficult, validating the cognitive load effect when the instructions are changed between two sets.

Participants made more commission errors in high cognitive load situations when faced with food stimuli. This was not the case for neutral stimuli, seeing as the proportion of errors for object pictures did not differ between the high cognitive load and the low cognitive load condition. This led us to conclude that cognitive load modulates inhibitory control, but only toward foods. The increase in mental effort that was required to process the instructions led participants to make significantly more impulsive detections, resulting in more commission errors. We can deduce that significant cognitive resources were needed for the integration and automatization of the new instructions. In the meantime, the amount of cognitive resources needed to inhibit the approach tendency elicited by HED foods was increased by the higher cognitive load. There were thus not enough resources allocated to inhibit interferences from prepotent responses, triggering commission errors. Indeed, the cognitive load effect indicates that there is a cognitive deficit in inhibitory control prior to behavioural disinhibition, as indicated by commission errors. This result correlates with previous research investigating the role of cognitive load in inhibitory control [84] and showing that working memory load (resulting here from the new set of instructions) interacts heavily with inhibitory control [28].

Food stimuli are salient, which induces an approach bias that interferes with the initiation of goal-directed behaviour on a cognitive level, leading to cognitive and behavioural deficits in inhibitory control. This process occurs in individuals regardless of weight status, and its intensity seems to vary in function of food characteristics (i.e. category and/or energy density). Moreover, the deficit in inhibitory control induced by food stimuli is modulated by the cognitive load in working memory, which means that the more mental effort the individual has to make while performing a task, the fewer resources are available to inhibit prepotent responses. This phenomenon leads to more disinhibition, meaning that individuals may be more likely to eat more HED foods when their cognitive load is heavier.

### Limitations

As discussed above, our study presents some limitations. First, we question the use of fruit stimuli as LED food stimuli. Indeed, fruits are frequently consumed in non-processed and raw forms, making it easier to distinguish them from objects than HED foods in the earliest stages of feature perception. Some empirical data from electroencephalography demonstrated that fruits do indeed undergo earlier processing. The pattern of evoked potentials (EPs) for the fruity quality of food stimuli seems distinct from the patterns of EPs observed for sweetness/saltiness and low/high energetic value [76]. Moreover, there is less diversity in the presentation of fruit in everyday life when compared with sweet HED foods (chocolate bars, cakes and pastries), which come in a variety of forms. In terms of perception, the distinction between raw and transformed food goes beyond the calorie content [85]. We hypothesize that identifying pictures of fruit over a short time during a single presentation might thus be facilitated because fruits are well-known and belong to a universal category [86]. There are limited options in the pairing of fruits to comparable HED foods because it is difficult to find sweet calorie-dense foods that are not processed and that belong to a universal category. In our study, we only used sweet stimuli for odour-congruency and literature fidelity reasons, but this remark may or may not refer to vegetables, which are also consumed raw and non-processed, but do not benefit from early perception facilitation [76]. There is a need to find pictorial LED stimuli that fit HED stimuli in visual and hedonic properties, but also in their intrinsic features such as degree of processing and distance from categorical prototype.

Several studies have observed interesting priming effects with the pear and pound cake odour, which are odorant mixtures [19,20,23]. These effects were observed in relation to weight status, which indicates the need to identify olfactory components that tap into specific (and unknown to date) mental representations contributing to weight-status specific responses. Concerning the implicit priming, we suggest that a context of more incentive facilitation (involving a less implicit sensory modality than olfaction, or in multi-modal priming) might have a stronger influence on top-down processing. However, we insist on using implicit priming to experimentally manipulate the effects of incidental cues from the environment in laboratory experiments seeing as non-attentively perceived cues appear to have a stronger effect on cognitive processing [23] and behaviour [39] than explicitly primed cues. Also, they are more reflective of the influences of environmental cues which often occur out of the individual’s attentional focus [7].

Moreover, we suppose that the different stimuli types elicited different attentional control patterns, with HED food stimuli more likely to attract attention, thus impairing attentional control. Unfortunately, our experiment was not designed to identify the phenomenon of attentional control toward foods, and reaction times do not represent a pure measure of distinct attentional mechanisms [33]. Such measures should be included in further experiments in order to refine our understanding of the role of the attentional functions in food stimuli processing, for instance by adding eye-tracking measurements into the experimental design, similar to the method tested by Doolan et al [87].

### Perspectives

#### Cognitive load in obesity

In the Ironic Process Theory [88], the daily life stressors increase cognitive load, which modulates inhibitory control. These synergic effects tend to produce behaviours opposite to what was primarily intended by the individual. Considerable research has shown that individuals with obesity and overweight are more at risk of exposure to daily life stressors: low income [89], anxiety [90], psychological health impairments [91], physical comorbidities [92], and discrimination and stigmatization in relation to body weight [93,94]. Considering all these aspects leads us to suppose that individuals with obesity might be subject to higher cognitive loads during daily decision-making, which could alter their inhibitory control and consequently, produce goal-unrelated behaviours. In our study, individuals were experimentally confronted to the same amount of cognitive load, which made it impossible to discriminate individual levels of inhibitory control toward foods according to weight status. We now suggest that variations in everyday cognitive load might explain some of the relationships between behaviourally reflected lack of inhibitory control facing foods and weight status that was identified in other studies. In future research, these relationships should be characterized in order to better understand overweight and obesity.

#### Implicit priming as a context of facilitation

Several studies focusing on inhibitory control manipulated the cognitive processing of food stimuli by creating a context of facilitation with priming (priming concepts of impulsivity [37] and unrestrained food consumption [38]), which led to interesting results. Nevertheless, such priming was explicit and is therefore not reflective of incidental food cues from the environment, which was part of the objective of our study. Different forms of implicit priming could be used in future research in order to assess the effects of implicit food cues on inhibitory control or other top-down processes toward foods in a unimodal or multimodal manner. For instance, the combination of auditory and olfactory priming has already been suggested as a means to influence individual food choices [17]. In future research, this type of multimodal priming could be used as an experimental context of facilitation in order to elicit a lack of inhibitory control for food intake.

## Conclusion

Our study highlights common mechanisms relative to the top-down processing of foods, regardless of individual weight status. Food stimuli are salient, which induces an approach bias that interferes with the initiation of goal-directed behaviour on a cognitive level, leading to cognitive and behavioural deficits in inhibitory control. This process occurs in individuals regardless of weight status, and its intensity seems to vary in function of food characteristics (i.e. category and/or energy density). This deficit in inhibitory control induced by food stimuli is modulated by the cognitive load in working memory, which means that the more mental effort the individual has to make while performing a task, the fewer resources are available to inhibit prepotent responses. Future research should focus on weight status in relation to cognitive load in order to improve our understanding of unhealthy food choices in obesity. Our data support that implicit priming selectively modulates bottom-up processing. Specific priming effects of food cues by weight status were also characterized in bottom-up processing, which opens a new path for research on mental representations activated by food cues among the weight status continuum.

## Supporting information

Full data are available in the Supporting Information Files, see S1 Dataset.

## Acknowledgements

We would like to thank the society Psytest, for lending us the five TAP versions, and especially Mr Benjamin Steves for his informative help about the material. We also would like to thank Jacques Maratray for developing modified Affective Shifting Task and Suzanne Rankin for English proofreading. We also thank the Chemosens platform for their help with participants’ recruitment, as well as Maya Filhon for her technical help during experimental sessions.

S1 Dataset. Full data available.

## Footnotes

1 Such processes are referred to as “bottom-up” or stimulus-driven processes, meaning that data from the environment drive our perception of stimuli

2 The extent of which a stimulus is salient

