## Supplementary material for "Implicit food odour priming effects on reactivity and inhibitory control towards foods": Full data are available in the Supporting Information Files, see S1 Dataset.: ReadMe-Mas-PlosOne-F509-2020.txt

Mas , Brindisi, Chabanet, Chambaron (2020)
Article : Implicit food odour priming effects on reactivity and inhibitory control towards foods
Related file : data-Mas-PlosOne-F509-2020.csv

Variables Description : 
id              : participant code (1 letter + 3 numbers)
weight.status   : weight status of the participant
	          N = normal-weight
		  S = overweight
		  O = obesity
type            : image type
	          HED=high-energy dense food picture
		  LED : low-energy dense food picture
		  CTL : object picture
odour           : P=pear odour
	          Q=pound cake odour
		  R=control (no odour)
gng             : whether the trial is a go or no go trial
	          go= go trial
		  nogo= no-go trial
CL              : cognitive load
	          CL- : low cognitive load
		  CL+ : high cognitive load
RT 	        : reaction time (in milliseconds)
error 	        : 0=no comission error
	          1= comission error
omission        : 0=no omission error
	          1=omission error
		  NA=trial not eligible to an omission error
hunger.before   : hunger before session 
hunger.after    : hunger after session
sex 	        : F=female
	       	  M=male
age 	        : age of participant
edu.lvl         : educational level
	       	  Bac-- : <=14 years
		  Bac++ : > 14 years
TAP.flexibility : global cognitive performance, flexibility subtest score 
TAP.gonogo	: global cognitive performance, Go/no-Go subtest
